# Spatio-temporal modelling of the effect of selected environmental and land-use factors on species-rich calcareous grasslands: overgrazing and nitrogen deposition

**DOI:** 10.1101/2022.11.29.518308

**Authors:** Christian Damgaard

## Abstract

The abundance of sensitive plant species in calcareous grasslands are threatened by agricultural intensification with nutrient addition and increased livestock densities as well as by land abandonment. In order to quantify the effect of selected environmental and land-use factors on the observed variation and changes in the vegetation of calcareous grasslands, large-scale spatial and temporal pin-point plant cover monitoring data are fitted in a structural equation model. The analyzed vegetation data come from 100 Danish sites monitored over an eight year period. The important sources of measurement- and sampling uncertainties have been included using a hierarchical model structure. Furthermore, the measurement- and sampling uncertainties are separated from the process uncertainty, which is important when generating ecological predictions that may feed into local conservation management decisions. There were significant negative effects of grazing and nitrogen deposition on the change in cover of sensitive plant species. Whereas the negative effect of nitrogen deposition on the cover of sensitive species was expected, it was surprising that the model results suggest that the class of sensitive species may be subjected to overgrazing by the grazing regimes that are currently applied at protected Danish calcareous grasslands. The standardized regression coefficients suggest that the effects of both grazing and nitrogen deposition are relatively benign compared to the effects of soil type, soil pH and precipitation. The relatively large effect of precipitation is interesting, since it is predicted that the amount of precipitation and distribution over the season will change due to climate change, and the results suggest that climate change may lead to important species compositional changes in calcareous grasslands. The fitted model may be used to quantify the uncertainties when generating ecological forecasting and local adaptive management plans.

## Introduction

North-European calcareous grasslands are species-rich semi-natural habitats typically found on nutrient-poor, well-drained calcareous to neutral soils with periods of summer droughts and are key habitats for many species: herbs, grazing animals, butterflies, reptiles and many birds (Olmeda et al. 2019). Calcareous grasslands have low productivity and persist due to extensive grazing and mowing. They are threatened both by agricultural intensification with nutrient addition and overgrazing due to increased livestock densities as well as by land abandonment, where grasslands are overgrown by shrubs and trees (Olmeda et al. 2019; Petit and Berien 2006; Timmermann et al. 2015).

Atmospheric nitrogen deposition has been shown to be associated with changes in species composition, where small, light-demanding and winter-green or evergreen plant species with smaller seeds and scleromorphic leaves were selected against (Diekmann et al. 2014). In several habitat types, nitrogen deposition has been associated with a decrease in species richness, but this seems not to be case for calcareous grassland (Maskell et al. 2010), although there has been a general decrease in the species richness of habitat specialist (Diekmann et al. 2019) and insect pollinated plants (Ehlers et al. 2021).

Changes in precipitation and temperature patterns due to climate changes may cause modifications in interspecific plant competitive interactions and alterations in plant community compositions (Herben et al. 2003; Hopkins and Del Prado 2007), although the abundance of different lifeforms in a calcareous grassland was relatively resistant to long-term climate manipulations (Grime et al. 2008).

Invasive species have also been mentioned as a possible threat to calcareous grasslands (Olmeda et al. 2019). In Denmark, invasive species were found in 6% of randomly placed circles with a radius of 5 meters, although this occurrence probability of invasive species did not change significantly in the period 2004– 2014 (Damgaard et al. 2019a).

Overall, the conservation status of calcareous grasslands is unfavorable and deteriorating in most of the European habitat range, although the conservation status inside the Natura 2000 network is better than outside (Olmeda et al. 2019). In Denmark, the conservation status of calcareous grasslands is determined using a multitude of criteria of different indicators (vegetation height, cover of shrubs and trees, Ellenberg N and R, soil phosphorous, number of characteristic species), which all should be fulfilled before the conservation status may be classified as favorable (Damgaard et al. 2019b; Nygaard et al. 2014).

To reverse this negative trend in conservation status, it is crucial to understand the key ecological requirements, which may vary at national and local level, for the establishment of conservation measures to ensure a favorable conservation status of the habitat. Furthermore, site level conservation objectives are needed for establishing the necessary conservation measures required for the habitat types and species that motivate the site designation (Olmeda et al. 2019).

Most large-scale studies on calcareous grassland vegetation have been conducted using spatial regression methods (Ridding et al. 2020), where it is assumed that the observed spatial patterns are due to different realizations of the studied ecological processes, and temporal dynamics may be ignored. This is problematic, since the habitats are known to be subjected to large changes in land use and climate, and there may be important time-delay effects from an environmental change to the possible biotic response (e.g. Jackson et al. 2009; Jackson and Blois 2015; Jackson and Sax 2010; Kratz et al. 2003; Svenning and Sandel 2013). Consequently, to correctly identify and quantify the effect of different drivers on calcareous grassland vegetation, it is essential to study both spatial and temporal variation (Damgaard 2019a; Diekmann et al. 2019).

Here, spatial and temporal pin-point plant cover data are fitted to a spatio-temporal structural equation model (SEM) in order to understand and quantify the effect of environmental variables and land-use on the observed variation and changes in the vegetation of calcareous grasslands. Furthermore, latent geographic factors of the large-scale spatial variation will be estimated (Ovaskainen et al. 2016). The SEM has been fitted within a Bayesian hierarchical model structure using latent variables in order to model the effect of measurement- and sampling uncertainties. The use of a hierarchical model structure is important since it has been demonstrated that ignoring measurement- and sampling uncertainties may lead to model- and prediction bias (Damgaard 2020; Damgaard and Weiner 2021). Furthermore, it is an advantage when making ecological predictions to separate measurement- and sampling uncertainties from process uncertainty. The hierarchical SEM approach and the motivation for using it are explained in more detail in Damgaard (2019b), where a similar model was fitted to ecological monitoring data from wet heathlands, and more generally in Damgaard (2022a).

The aim of the present study is to model the effect of selected environmental and land-use variables; soil type, pH, nitrogen deposition, precipitation and grazing on the large-scale spatial distribution and temporal changes of the calcareous grassland vegetation. The obtained model results will be discussed in light of the above-mentioned threats to calcareous grassland habitats and their possible effect on the conservation status.

It is not possibly to model each species separately when modelling plant community dynamics, and therefore it was necessary to aggregate the plant species into species classes (Damgaard 2019b). This species aggregation is somewhat arbitrary, but has to reflect the aim of the study. Since the main aim of this study is to understand the effect of selected environmental and land-use variables on the conservation status of calcareous grasslands, I have chosen to aggregate the pin-point cover data of all higher plants into sensitive species, other graminoids and other forbs. The species belonging to the class “sensitive species” were assessed by experts to be sensitive or very sensitive towards excess soil nutrients, reduced water table or/and change in land-use (Fredshavn and Ejrnæs 2009; Nygaard et al. 2019). Consequently, we expect to find negative effect of e.g. nitrogen deposition on the class of sensitive species, but the importance of the effect is generally unknown. Many of the sensitive species are characteristic species of the habitat, and the presence of sensitive species is a positive indicator of a favorable conservation status.

## Materials and Methods

### Sampling design

Hierarchical time series data from 100 calcareous grassland sites (Fig. 1) that had been monitored at least three times in the period from 2007 to 2014 were used in the analysis. Sixty-six of the 100 sites are NATURA 2000 habitat sites and protected under the habitat directive (EU 1992). All sites included several randomly-positioned plots classified as calcareous grassland (EU habitat type: 6210) according to the habitat classification system used for the European Habitat Directive (EU 2003). The area of the sites ranged from 0.20 ha to 304 ha, with a median area of 9.8 ha. A total of 1174 unique plots were used in the analysis. The sampling intensity was irregular among sites and years, but typically between ten and forty plots were sampled from each site at each time point. Including resampling over the years, a total of 3535 plot data were used in the analyses. The plots were resampled with GPS-certainty (< 10 meters), and the analysis of the hierarchical data set were therefore performed at the site level.

**Fig. 1.**
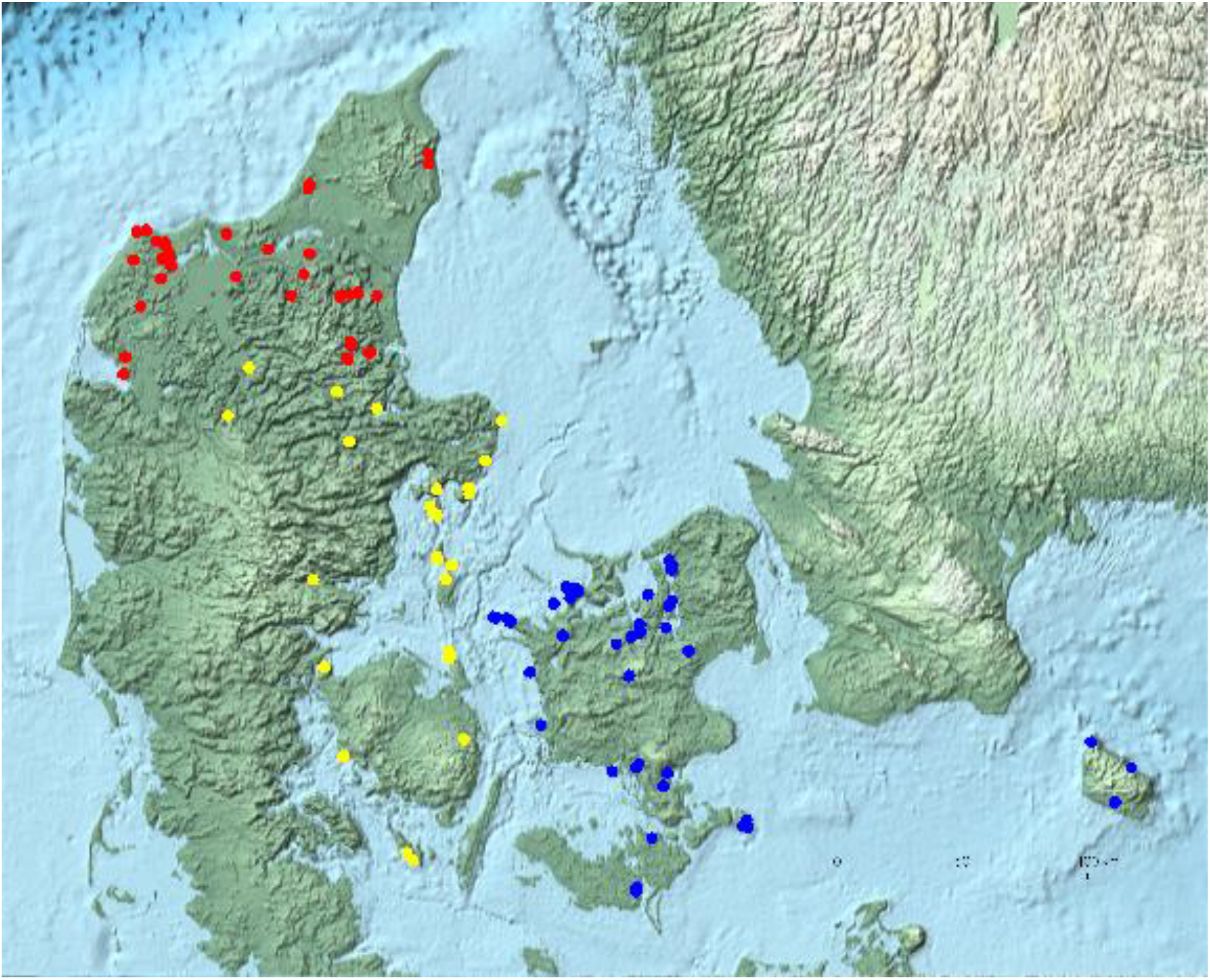
Map of the 100 Danish calcareous grassland sites. The different colors represent a classification of the different sites into three geographical regions.

### Variables and measurements

The used data are a subset of the data collected in the Danish habitat surveillance program NOVANA (Nielsen et al. 2012; Nygaard et al. 2016) and relevant climate data (DMI 2014). Both data sets are publically available.

#### Plant cover data

The plant cover, which is the relative projected area covered by a species, was measured for all higher plants by the pin-point method using a square frame (50 cm X 50 cm) of 16 grid points that were equally spaced by 10 cm (Nielsen et al. 2012). At each grid point, a thin pin was inserted into the vegetation and the plant species that was touched by the pin was recorded and used as an estimate of cover (Damgaard and Irvine 2019; Levy and Madden 1933; Lindquist 1931). Cover and trends in cover for the 50 most common species in Danish calcareous grassland are summarized in Table S1.

Since the pin-point cover data were recorded for each pin separately, the species cover data are readily aggregated into cover data for classes of species at a higher taxonomic or functional level. At each grid point the pin may hit different plant species from the same species class, and in those cases the hits are only counted as a single hit of the species class at the grid point.

In this study, the observed higher plant species were divided into three species classes: sensitive species, other graminoids and other forbs, where the class of sensitive species comprised 1058 plant taxa (Table S2). The class of sensitive species were defined as the species that scored four or above in a scoring system of sensitive species where seven is the maximum. The scoring system was developed by an expert evaluation of the sensitivity of the species towards environmental drivers and if the species were typically found at sites at high conservation status (Fredshavn and Ejrnæs 2007). The mean site covers of the three species classes are shown in Fig. 2.

**Fig. 2.**
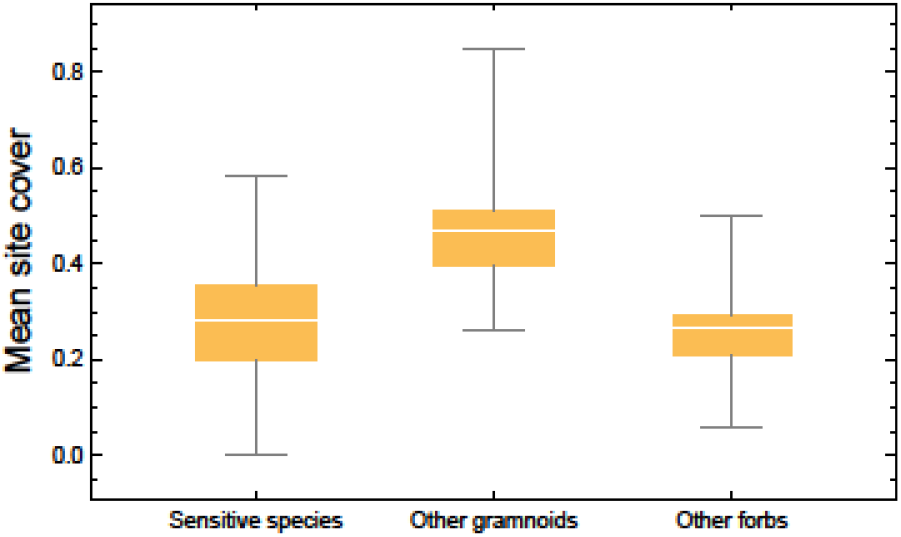
Box plots of the mean site cover of the three plant species classses.

The joint distribution of the multivariate cover data of the three species classes were analyzed using a reparametrized Dirichlet - multinomial distribution, where the typically observed spatial aggregation of plant species is taken into account (Damgaard 2015; 2018).

#### Nitrogen deposition

Nitrogen deposition at each plot was calculated for each year using a spatial atmospheric deposition model in the period from 2005 to 2014 (Ellermann et al. 2012). The mean site nitrogen deposition ranged from 5.66 kg N ha^−1^ year^−1^ to 17.53 kg N ha^−1^ year^−1^, with a mean deposition of 11.27 kg N ha^−1^ year^−1^. Anthropogenic nitrogen deposition has reached a maximum and is currently decreasing in Denmark (Ellermann et al. 2018).

#### pH

Soil pH was measured in randomly selected plots from the uppermost 5 cm of the soil (four samples were amassed into a single sample). The soil sampling intensity was irregular among sites and years, but typically between one and four plots were sampled from each site at each time point. When a plot was resampled, the pH at the plot was calculated as the mean of the samples. In total, 1215 soil pH measurements were used in the analysis. The soils were passed through a 2mm sieve to remove gravel and coarse plant material, and pH_KCl_ was measured on a 1 M KCl-soil paste (1:1). The measured soil pH ranged from 3.2 to 8.0, with a mean soil pH of 4.81. Note that pH_KCl_ values are generally one pH unit lower than if water was used as a buffer.

#### Soil type

The texture of the top soil for each site was obtained from a raster based map of Danish top soils (Greve et al. 2007). The categorical classification of the soil (JB-nr.) is made on an ordinal scale with decreasing particle size: 1: coarse sandy soil, 2: fine sand soil, 3: coarse loamy soil, 4: fine loamy soil, 5: course sandy clay soil, 6: fine sandy clay soil, 7: clay soil, 8: thick clay soil. The mean soil type was 3.94.

#### Precipitation

Site-specific precipitation was measured by the average annual precipitation in the period 2001 to 2010, with a spatial resolution of 10 km (DMI 2014). The annual precipitation ranged from 568 mm to 858 mm, with a mean precipitation of 697 mm.

#### Grazing

Land-use was summarized by possible signs of grazing, e.g. the presence of livestock or short vegetation within fences was recorded by the observer at each plot for each sampling year since 2007 as a binary variable (sign of grazing = 1, no sign of grazing = 0), i.e. if grazing was 0.5 then this probability may arise by a number of ways e.g. if half the plots at the site showed signs of being grazed each year or all plots were grazed every second year. The mean grazing probability at the site level was 0.54. The grazing variable does not include information on which animals were used for grazing, stocking densities or grazing duration, and is therefore a quite imprecise variable that has to be interpreted together with general knowledge on the typically used grazing regime in Danish calcareous grassland (Danish Nature Agency 2014).

Only a few sites were mown and the mean mowing probability at the site level was only 2%. Consequently, mowing was not considered in this analysis.

#### Geographic regions

The 100 calcareous grassland sites were grouped into three geographic regions (Fig. 1) that were used to investigate possible latent geographic factors (Ovaskainen et al. 2016).

### Structural equation model

The variables were modelled in a SEM based on current qualitative knowledge on the causal effect relationships among the studied abiotic variables and grazing and their effect on the vegetation in calcareous grasslands (Olmeda et al. 2019). The SEM was fitted within a Bayesian hierarchical framework with structural equations and measurement equations in an acyclic directed graph (Fig. 3), where each arrow is modelled by a conditional likelihood function and uninformative prior probability distributions (Damgaard 2019b).

**Fig. 3.**
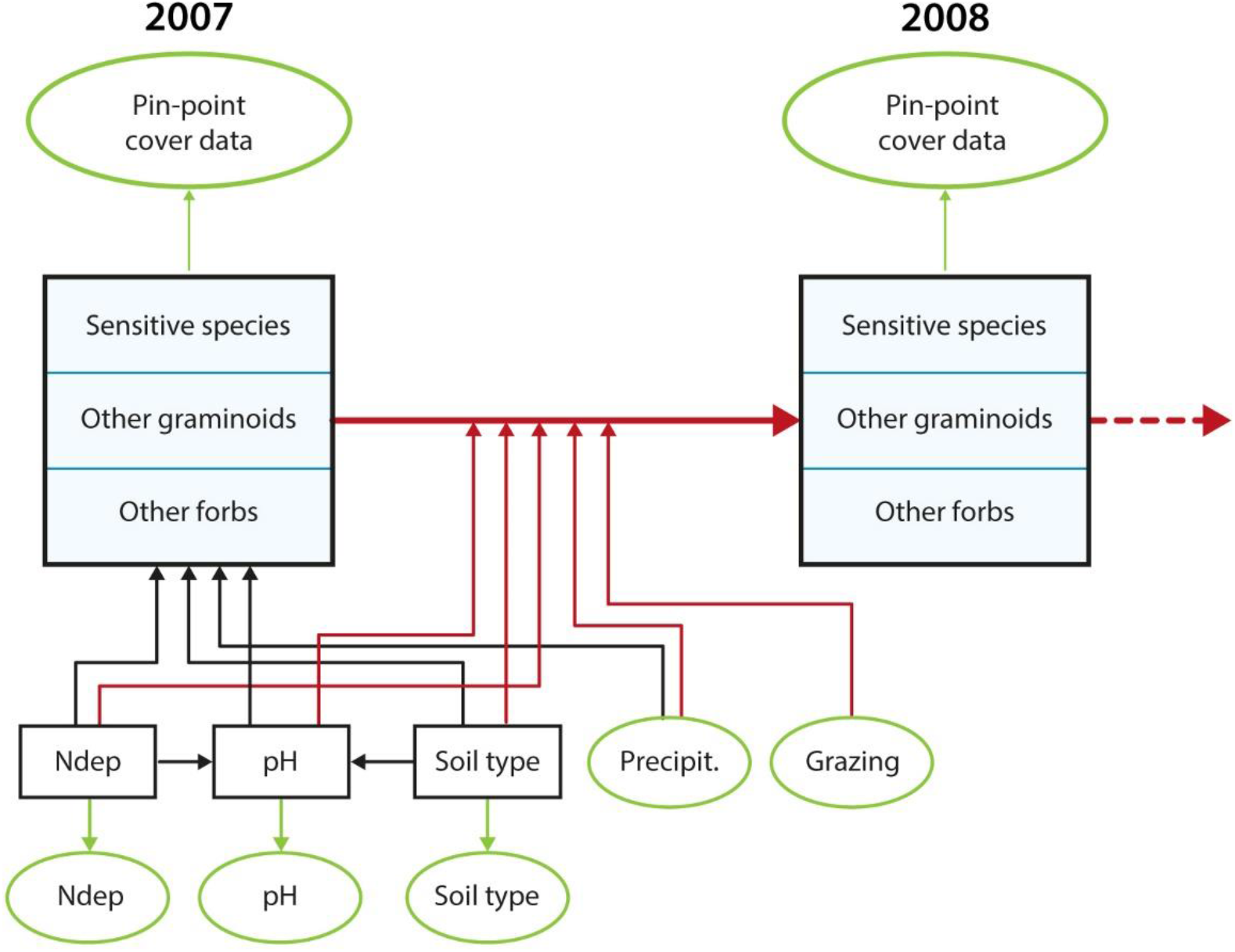
Outline of SEM. The spatial variation in vegetation cover in 2007 is modelled by nitrogen deposition (Ndep), soil pH (pH), soil type and precipitation (Precipit.). The yearly change in vegetation cover from 2007 to 2014 (only a single yearly change is shown in the figure) is modelled by all the former variables as well as grazing. The black boxes are latent variables and the green ovals are data. The black arrows denote spatial processes, and the red arrows denote temporal processes.

#### Structural equations

The structural equations consist of a mixture of measured data and latent variables, which are the true, but unknown, mean of the underlying data (Fig. 3). The latent variables are denoted by upper case Latin letters, data are denoted by lower case Latin letters, and model parameters are denoted by Greek letters.

The relative cover of the three species classes, *s*: sensitive species, other graminoids, and other forbs at each site, *i*, for the first year (1 = 2007) and for all subsequent years, *y*, where cover was measured, are modeled as latent variables, *Q*_*s*,*i*,*y*_. Furthermore, the latent geographic factor (*Geo*_*i*_) and the true, but unknown, mean of nitrogen deposition (*N*_*i*_), soil pH (*R*_*i*_) and soil type (*S*_*i*_) were modeled by site-specific latent variables. Site-specific average precipitation is denoted by *p*_*i*_.

The large-scale spatial variation in the relative cover the first year is modelled as, 

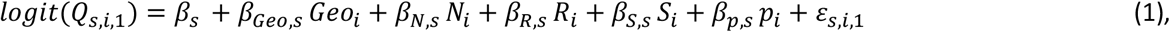

where the structural uncertainty in the spatial variation of the logit-transformed relative covers are assumed to be normally distributed, *ε*_*s*,*i*,1_~*N*(0, *σ*_*S*,*s*_).

The yearly change in the logit-transformed relative cover in subsequent years is modelled as, 
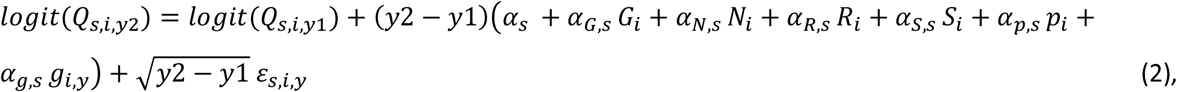

where (*y*2 − *y*1) is the years between successive sampling at site *i*, *G*_*i*_ is the latent variable that measures grazing, and the structural uncertainty in the temporal variation of the logit-transformed relative covers are assumed to be normally distributed, *ε*_*s*,*i*,*y*_~*N*(0, *σ*_*T*,*s*_).

The site-specific soil pH in the top soil was modelled as, 
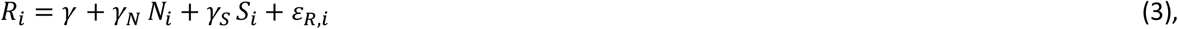

where the structural uncertainty in soil pH is assumed to be normally distributed, *ε*_*R*,*i*_~*N*(0, *σ*_*SR*_).

The latent geographic factors were assumed to come from a Gaussian process, *G*~*N*(0,1) (Haran 2011).

#### Measurement equations

The measurement equations link the latent variables to the measured or model-calculated data with conditional likelihood functions (Fig. 3):

The observed aggregated pin-point cover data of sensitive species, other graminoids, and other forbs at site *i*, plot *j* and year y, *q*_*s*,*i*,*j*,*y*_ were assumed to be jointly distributed according to a Dirichlet - multinomial mixture distribution, *q*_*s*,*i*,*j*,*y*_~*DM*(*Q*_*s*,*i*,*y*_, *δ*), with *Q*_*s*,*i*,*y*_ as the relative mean cover of sensitive species and other graminoids at site *i*, and small-scale spatial aggregation parameter *δ* (Damgaard 2015; 2018). If the plot had been sampled before then within-plot auto-correlation was taken into account by, *q*_*s*,*i*,*j*,*y*_~*DM*(*Q*_*s*,*i*,*y*_ + *ρ*_*s*_(*q*_*s*,*i*,*j*,*yp*_ − *Q*_*s*,*i*,*yp*_), *δ*), where *ρ*_*s*_ are correlation coefficients and *yp* denote the year of the pervious sample at the plot (Damgaard 2012).

The average model calculated nitrogen deposition at site *i* and year *y*, *n*_*i*,*y*_, was assumed to be normally distributed, *n*_*i*,*y*_~*N*(*N*_*i*_, *σ*_*N*_), where *σ*_*N*_ is the yearly variation in model calculated nitrogen deposition.

The measured soil pH at site *i* and plot *j*, *r*_*i*,*j*_, was assumed to be normally distributed, *r*_*i*,*j*_~*N*(*R*_*i*_, *σ*_*R*_), where *σ*_*R*_ is the within site variation in soil pH. At three sites there were no measurements of soil pH, and at these sites *R*_*i*_was not conditioned by measured data, but only by the structure of the fitted SEM (Fig. 3).

The predicted texture of the top soil at site *i* and plot *j*, *s*_*i*,*j*_ was assumed to be normally distributed, *s*_*i*,*j*_~*N*(*S*_*i*_, *σ*_*S*_), where *σ*_*S*_ is the within site variation in soil type.

### Estimation and statistical inference

The SEM was parameterized using numerical Bayesian methods. The overall likelihood function of the SEM model was found by multiplying the conditional likelihood functions specified above and outlined in figure 3, using a first order Markov assumption (Clark 2007). The joint posterior probability distribution of the parameters and the latent variables were calculated using Markov Chain Monte Carlo (MCMC) and Metropolis-Hastings, simulations with a multivariate normal candidate distribution. The MCMC had 200,000 iterations, with a burn-in period of 140,000 iterations. The fitting of the hierarchical SEM is a rather slow process and the MCMC simulations were running for 18 months on a workstation computer.

The prior probability distributions of all parameters were assumed to be uniformly distributed either as improper priors or in their specified domain, except standard deviation parameters of the normal distribution that were assumed to be inverse gamma distributed, (*σ*_*x*_~*IG*(0.1,0.1). The prior probability distributions of the cover latent variables were assumed to be uniformly distributed within 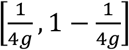, where *g* is the number of pins in a pin-point frame (Damgaard 2012).

Plots of the sampling chains of all parameters and latent variables were inspected to check the mixing properties of the used sampling procedure. Additionally, the overall fitting properties of the model were checked by inspecting the regularity and shape of the marginal distribution of parameters as well as the distribution of the deviance (= −2 *log L*(*Y*|*θ*)). The efficiency of the MCMC procedure was assessed by inspecting the evolution in the deviance.

In order to check the fit of the model, the mean latent logit-transformed cover variables were plotted against the expected logit-transformed cover for both the spatial and temporal structural processes. Furthermore, the Dunn–Smyth residuals (Dunn and Smyth 1996) of the marginal observed cover data of the three species classes were calculated with the mean latent cover variables as the expected covers (*q*_*i*_) and the mean small-scale spatial aggregation parameter (*δ*) (Damgaard 2018).

Statistical inferences on the parameters were based on the estimated marginal posterior probability distribution of the parameters. Standardized regression coefficients (path coefficients) were calculated by multiplying the estimated partial regression coefficients with the standard deviation of the independent latent variable and dividing with the standard deviation of the dependent latent variable.

All calculations were done using *Mathematica* version 11.0.1 (Wolfram 2016), and the *Mathematica* notebook with additional diagnostic plots is provided as an electronic supplement.

## Results

The studied abiotic and land-use variables covaried at the 100 calcareous grassland sites (Fig. S1), which again is expected to lead to covariance among parameter estimates (Table S3) and affect the fitting properties of the model negatively. Plots of the mean latent vs. expected logit-transformed cover variables demonstrated a relatively poor fit of the large-scale spatial variation in cover (Fig. 4A; 38% of the variation is explained). However, the model fitted the temporal process of the change in cover very well (Fig. 4B; >99% of the variation is explained). Generally, the distribution of the Dunn–Smyth residuals of the marginal observed cover data of the three species classes were approximately normally distributed (Fig. S2).

**Fig 4.**
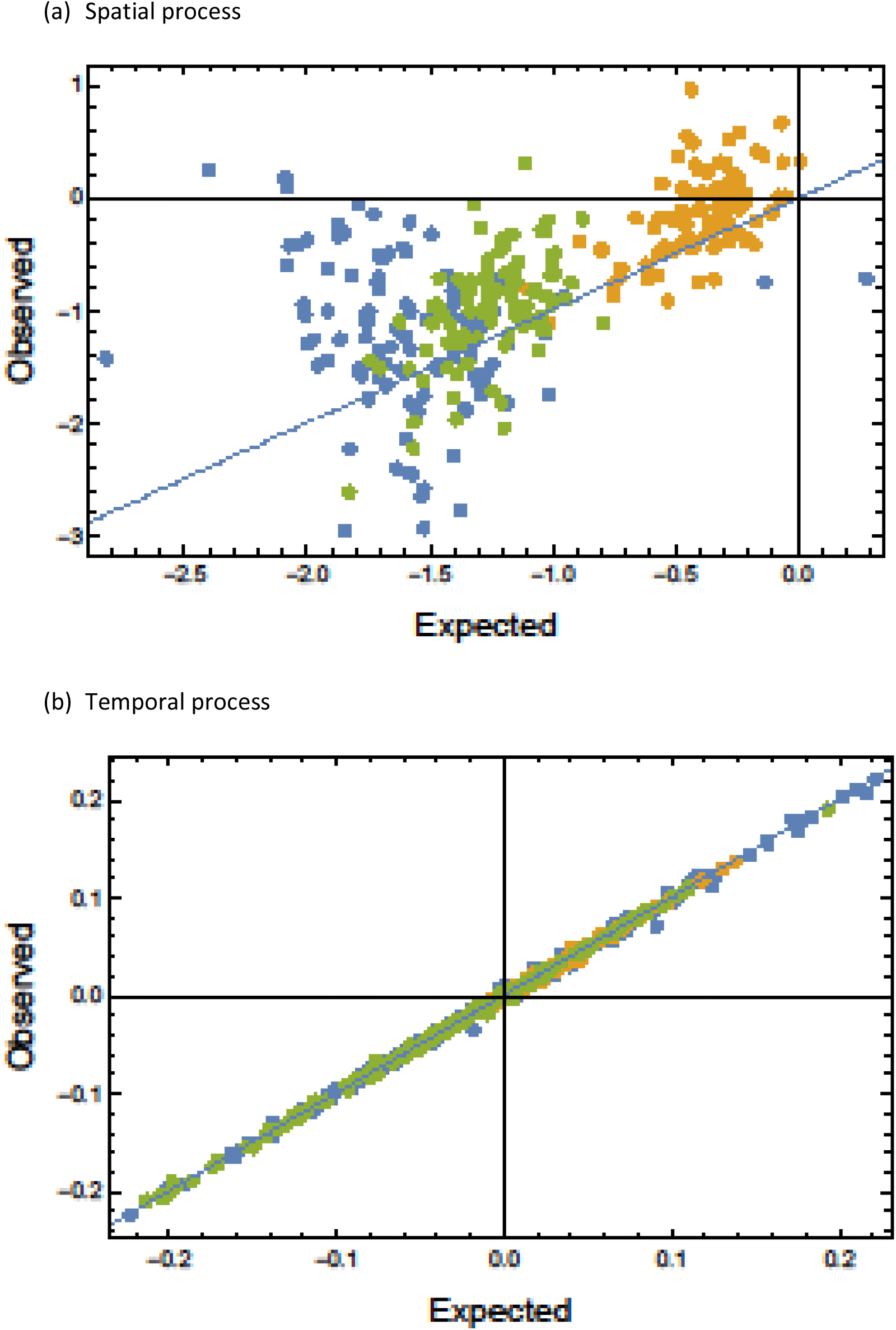
Plots of observed vs. expected of initial logit-transformed plant cover (a) and subsequent change of logit-transformed cover (b). Blue: sensitive species, yellow: other graminoids, green: other forbs.

To prevent possible prediction bias, the different sources of uncertainty, i.e. measurement – and sampling uncertainty when measuring plant cover, nitrogen deposition, soil pH and soil type, as well as, structural uncertainties due to the modelled soil pH, large-scale spatial variation and the temporal processes, were modelled explicitly. The most important source of measurement uncertainty was the plant cover measurement due to the significant small-scale spatial aggregation of plant species, which was modelled by the parameter *δ* in the Dirichlet - multinomial mixture distribution. The median estimated value of *δ* was 0.10 with a relatively narrow credible interval (Table 1). Generally, the structural uncertainties are low, except for the large-scale spatial variation, which is high (*σ_S_*, Table 1, Fig. 4a).

**Table 1.**
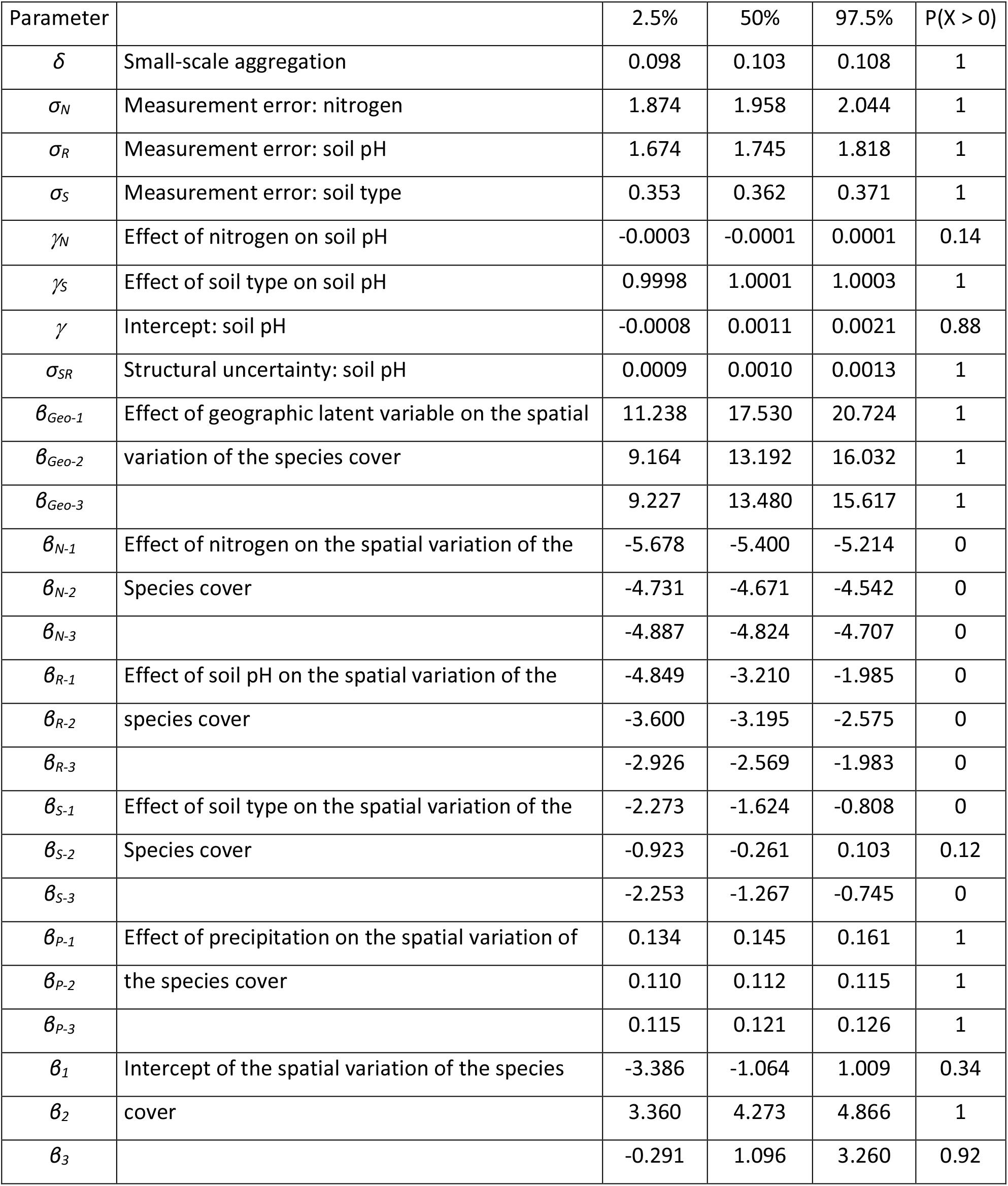
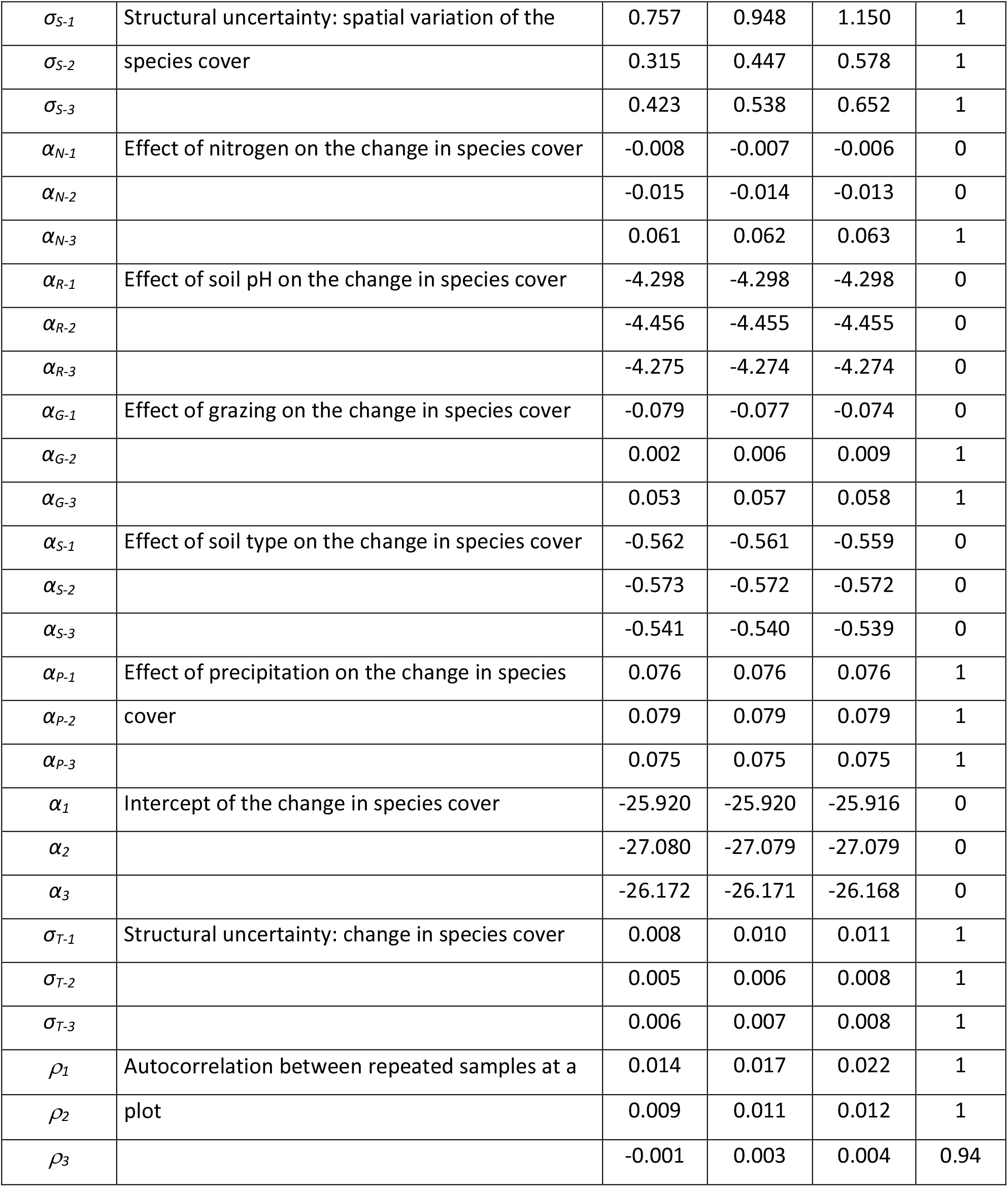
Marginal posterior probability distribution of SEM parameters shown by the 2.5%, 50% and 97.5% percentiles as well as the probability that the parameter is larger than zero. The plant species are denoted by numbers, 1: Sensitive species, 2: Other graminoids, 3: Other forbs.

Most of the regression parameters that measure the effect of the abiotic variables on the vegetation were significantly different from zero (Table 1), suggesting that the studied environmental and land-use factors have a regulating effect on both the large-scale spatial variation in cover of the three species classes as well as local plant community dynamics in calcareous grasslands.

Where was considerable large-scale spatial variation in cover of the three species classes (Table 1, Fig. 5A), but since the model only explained 38% of the variation and each of the abiotic variables had the same qualitatively effect on all three species classes, no clear pattern of the large-scale spatial variation can be extracted from the model.

**Fig 5.**
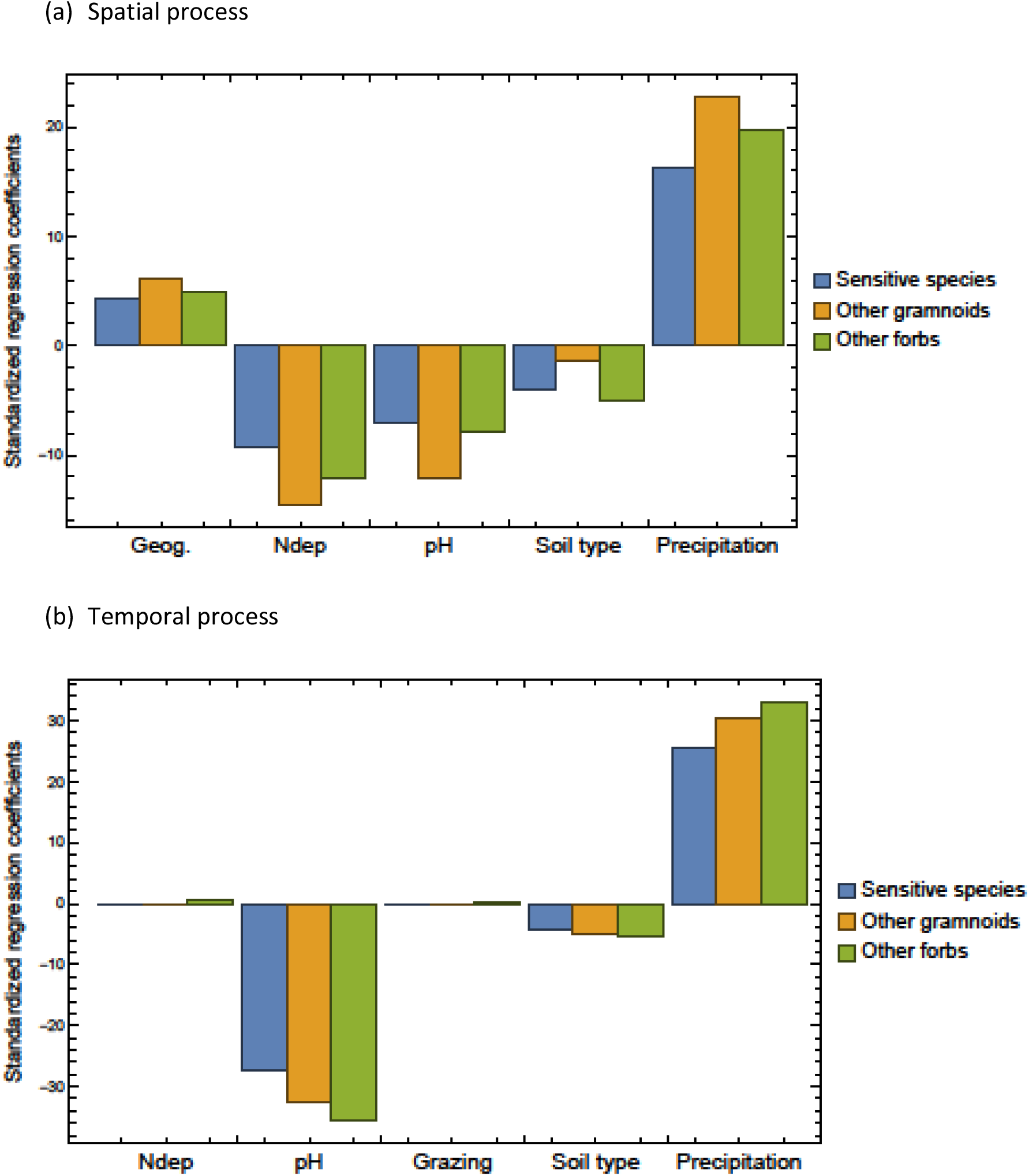
Standardized regression coefficients of the SEM for the initial spatial effects (a) and the later temporal effects (b).

The fit of the temporal process model was very good, and >99% of the observed variation in the change in cover during the study period was explained by the model. This high level of model fit is surprising and is higher than when the model has been used before on other habitat types (Damgaard 2019b; 2022b), although in those cases the model was fitted to four species classes, whereas here it was only fitted to three species classes. Increasing nitrogen deposition had a significant negative effect on the cover of sensitive species and other graminoids and a significant positive effect on other forbs. Increasing soil pH had a significant negative effect on the cover of all species classes. Grazing had a significant negative effect on sensitive species and significant positive effects on other species. Increasing the clay content of the soil had a significant negative effect on the cover of all species classes. Increasing precipitation had a significant positive effect on the cover of all species classes (Table 1). When regarding the standardized regression coefficients of the change in cover, the most important factors were soil pH and precipitation followed by soil type (Fig. 5B).

There were no significant effects of nitrogen deposition on soil pH (*γ_N_*, Table 1), but soil pH was found to be significantly higher on more clayey soils compared to sandy soils (*γ_S_*, Table 1). It was therefore concluded that the effect of soil type on the vegetation encompassed both direct effects of soil type and indirect effects mediated by the effect of soil type on soil pH. On the other hand, the effect of nitrogen deposition was mainly through direct effects on the vegetation and not by possible soil acidification.

## Discussion

There were significant negative effects of grazing and nitrogen deposition on cover of sensitive plant species. Whereas the negative effect of nitrogen deposition on the cover of sensitive species was expected, it was surprising that the model results suggest that the class of sensitive species may be subjected to overgrazing by the grazing regimes that are currently applied at Danish calcareous grasslands (although see Olmeda et al. 2019; Petit and Berien 2006). Especially since 66 of the 100 sites are NATURA 2000 habitat sites and protected under the habitat directive (EU 1992) and are expected to be subjected to adequate conservation management practices. Experiments with year-round low intensity grazing regimes have demonstrated an increase in the species richness of habitat characteristic species (Köhler et al. 2016), and this may be a promising avenue for future conservation management of calcareous grasslands.

The standardized regression coefficients suggest that the effects of both grazing and nitrogen deposition are relatively benign compared to the effects of increasing clay content, increasing soil pH and increasing precipitation. Of these, the relatively large effect of precipitation is interesting, since it is predicted that the amount of precipitation and distribution over the seasons will change due to climate change. In Denmark, the annual precipitation is predicted to increase, but with decreasing summer precipitation and longer summer drought periods (DMI 2017). Based on the current model results, it is not possible to predict which of the three species classes will be favored as a consequence of climate change, but it may be concluded that climate change may lead to significant species compositional changes in calcareous grasslands.

The used grazing data in this study are far from perfect; in a sampling year, it was assessed for each plot whether there were any signs of livestock grazing, and this assessment was recorded as a binary yes or no variable. Consequently, we have no information on the type of livestock, stocking densities or duration. However, it is assumed that there is a typical grazing system to manage calcareous grassland, and this grazing system has some variation in type of livestock, stocking densities or duration (Danish Nature Agency 2014). The results of this study indicate that this typical grazing system may be too intensive, since the cover of the aggregated class of sensitive species decreases at sites where grazing is recorded. This conclusion is in agreement with a European scale risk assessment of calcareous grasslands by Petit and Elbersen (2006), who found that 40% of the habitat area had a high or very high risk of being overgrazed, with a stocking density well above the recommended density of 0.25 units ha^−1^.

In order for the ecosystem model to be operational, the calcareous grassland vegetation has to be aggregated into a relatively few number of species classes. Here it was chosen to aggregate the species into only three species classes, where the class of sensitive species is directly relevant for the conservation of calcareous grassland habitats, and the relative abundance of the classes of *other graminoids* and *other forbs* may be used as an indicator for the conservation status of habitats. However, if the local nature management targets specific dominant and problematic plant species (Koerner et al. 2018; Pecháčková et al. 2010), then more detailed autecological information of dominant plant species may complement the model results.

The study corroborates the earlier findings that nitrogen deposition affects the species composition of calcareous grasslands (Diekmann et al. 2014; Ridding et al. 2020). The observed changes in species composition in Danish calcareous grassland sites with a mean nitrogen deposition of 11.3 kg N ha^−1^ year^−1^ may be compared with the estimated empirical critical load of nitrogen deposition in calcareous grasslands that is between 10 – 20 kg N ha^−1^ year^−1^ (Bobbink et al. 2022) and the biodiversity based critical load for the habitat type between 5 – 7 kg N ha^−1^ year^−1^ (Bak 2014).

It is generally expected that nitrogen deposition may lead to increased acidification (Bobbink et al. 2010; Williams and Anderson 1999). However, in Danish calcareous grasslands there was no significant large-scale spatial association between nitrogen deposition and soil pH, and this observed decoupling of nitrogen deposition and soil pH is in agreement with earlier findings in wet heathlands (Damgaard 2019b; Damgaard et al. 2014) and acid grasslands (Damgaard 2022b). The potential soil acidification effects of nitrogen deposition are expected to be caused by either nitrate leaching or removal of base cations (Williams and Anderson 1999), and the earlier and present results suggest that both soil process are limited in Danish light-open habitats.

Soil type and soil acidity are known to be important environmental factors that determine the functional and realized niche of plant species, where less acidic soils have a higher species richness (Ellenberg 1979; Pärtel 2002). As expected, relatively sandy soils were observed to be relatively more acidic, and the estimated effect of soil type on the vegetation included both direct effects of soil type and indirect effects mediated by the effect of soil type on soil pH.

In this study the most important sources of measurement and sampling uncertainties are explicitly included in the model and the measurement and sampling uncertainties are separated from the process uncertainties. These model features are essential for avoiding model- and prediction bias (Damgaard 2020; Damgaard and Weiner 2021), and possibly necessary when using ecosystem models for generating credible ecological predictions (Damgaard 2022a) and local adaptive management plans (Damgaard 2021).

## Acknowledgements

There were no conflicts of interest.

CD has analyzed the data and written the paper.

## Data availability

The used data are all in public domain, e.g. at https://naturdata.miljoeportal.dk/.

## Electronic supplements

The electronic supplements may be found at DOI 10.17605/OSF.IO/T58PS

Table S1: Cover data.

Table S2: Sensitive species.

Fig S1. Pairwise scatter plot of the mean at site level for all variables. Table S3: Correlation matrix of parameters.

Fig. S2. Dunn–Smyth residuals of the marginal observed cover data of the three species classes.

Mathematica notebook

